# Replicability of time-varying connectivity patterns in large resting state fMRI samples

**DOI:** 10.1101/172866

**Authors:** Anees Abrol, Eswar Damaraju, Robyn L. Miller, Julia M. Stephen, Eric D. Claus, Andrew R. Mayer, Vince D. Calhoun

**Affiliations:** The Mind Research Network, Albuquerque, New Mexico, USA.; Department of Electrical and Computer Engineering, University of New Mexico, Albuquerque, New Mexico, USA

## Abstract

The past few years have seen an emergence of approaches that leverage temporal changes in whole-brain patterns of functional connectivity (the chronnectome). In this chronnectome study, we investigate the replicability of the human brain's inter-regional coupling dynamics during rest by evaluating two different dynamic functional network connectivity (dFNC) analysis frameworks using 7500 functional magnetic resonance imaging (fMRI) datasets. To quantify the extent to which the emergent functional connectivity (FC) patterns are reproducible, we characterize the temporal dynamics by deriving several summary measures across multiple large, independent age-matched samples. Reproducibility was demonstrated through the existence of basic connectivity patterns (FC states) amidst an ensemble of inter-regional connections. Furthermore, application of the methods to conservatively configured surrogate datasets establishes that the correlation structures in the data do not arise by chance. This extensive testing of reproducibility of similarity statistics also suggests that the estimated FC states are robust against variation in data quality, analysis, grouping, and decomposition methods. We conclude that future investigations probing the functional and neurophysiological relevance of time-varying connectivity assume critical importance.

**Highlights:**

- Replicability in dynamic functional connectivity state measures was investigated.
- Twenty-eight samples each with two hundred and fifty rest-fMRI datasets were studied.
- State profiles were modelled using two (clustering and fuzzy meta-state) approaches.
- Both approaches showed high consistency for a range of model orders.
- Surrogate testing confirmed state summary measures to be statistically significant.

## 1. INTRODUCTION

More than two decades ago, the pivotal discovery of intrinsic low frequency fluctuations of blood flow and oxygenation levels in the functional magnetic resonance imaging (fMRI) modality was heralded as a neurophysiological index, and more specifically a manifestation of the intrinsic functional connectivity (FC) of the human brain (Biswal, Yetkin et al. 1995). This finding facilitated the study of FC in the neuroimaging research community, and since then numerous studies investigating characterization of partial-brain or whole-brain interactions stimulated by specific tasks or merely from spontaneous resting state activity have laid the foundation of our basic understanding of FC in the healthy and the diseased human brain. In an effort to boost discovery science in the functional neuroimaging community, traditional hypothesis-driven task-based studies paved the way for resting state fMRI approaches. Despite the unconstrained nature of the resting state experiments, the distributed networks or signal variations exhibiting temporal correlation in resting state fMRI decompositions, referred to as resting state networks (RSNs), were proven to have high levels of reproducibility thus suggesting a common architecture for the functional connectome (Damoiseaux, Rombouts et al. 2006, Fox and Raichle 2007, Margulies, Kelly et al. 2007, Shehzad, Kelly et al. 2009, Smith, Fox et al. 2009, Van Dijk, Hedden et al. 2010, Zuo, Kelly et al. 2010, Dansereau 2016). Additionally, consistency in the baseline functional activity of the brain (Damoiseaux, Rombouts et al. 2006) and reliability in some rest-fMRI measurements (Zuo and Xing 2014) has been suggested. Although there is a low degree of consensus on linkage of fMRI fluctuations with neural activity, there is rapidly growing literature providing evidence of association of the decomposed ICNs to underlying neuronal connectivity (Mantini, Perrucci et al. 2007, He, Snyder et al. 2008, Shmuel and Leopold 2008, Britz, Van De Ville et al. 2010, de Pasquale, Della Penna et al. 2010) which motivates investigations of spontaneous FC with great optimism.

Studies assessing FC primarily leverage seed-based correlation analysis (SCA) and spatial independent component analysis (ICA) to decompose brain signals into distributed networks exhibiting high temporal correlation in intrinsic activity (Joel, Caffo et al. 2011). SCA decompositions feature computation of pairwise correlation in time-courses corresponding to the predefined brain regions of interest (Biswal, Yetkin et al. 1995, Fox, Snyder et al. 2005). Several pre-defined ROI atlases such as the automated anatomical labeling atlas (Tzourio-Mazoyer, Landeau et al. 2002), the Talairach and Tournoux atlas (Lancaster, Woldorff et al. 2000), the Eickhoff-Zilles (Eickhoff, Stephan et al. 2005), the Harvard-Oxford atlas (Makris, Goldstein et al. 2006), and the Craddock atlas (Craddock, James et al. 2012) are leveraged to set up seeds for which pairwise functional connectivity is estimated. These seed based approaches have been largely successful in revealing useful information on brain-wide FC. Another widely used method in estimating seeds is the spatial ICA decomposition method (McKeown, Makeig et al. 1998, McKeown and Sejnowski 1998) that allows for measurement of network connectivity in multiple data-driven regions of interest. This method yields consistent spatially segregated and functionally homogeneous RSNs by exploiting independence in the spatial domain as opposed to assessment of fixed brain voxels. Consequently, current multi-subject studies frequently make use of the group ICA (gICA) technique (Calhoun, Adali et al. 2001) for extracting brain networks while retaining individual subject variability (Erhardt, Rachakonda et al. 2011, Allen, Erhardt et al. 2012).

More recently, there has been a major paradigm shift from measuring the whole-brain FC in the form of time averaged connectivity metrics (e.g. correlation, coherence, mutual information, etc.) to additional exploration of the time-varying (referred to as non-stationary, non-static or dynamic in previous literature) nature of the underlying fluctuations (Chang and Glover 2010, Musso, Brinkmeyer et al. 2010, Sakoglu, Pearlson et al. 2010, Liu, Zhu et al. 2011, Cribben, Haraldsdottir et al. 2012, Yuan, Zotev et al. 2012, Hutchison, Womelsdorf et al. 2013, Lindquist, Xu et al. 2014, Leonardi and Van De Ville 2015) in both task based and resting state fMRI. Notably, such studies evaluate time-varying FC between different brain regions (second order time-varying statistics) as compared to studying the coherent instantaneous fluctuations in the fMRI signal (first order time-varying statistics) that have been shown to account for variability in task-based BOLD responses as extensively reviewed in Fox and Raichle (2007). Similar to the static resting state FC literature, the ultimate objective of the time-varying resting state FC studies is to understand the driving mechanisms and cognitive implications of the observed fluctuations in FC of the brain regions. Studies (Chang, Liu et al. 2013, Tagliazucchi and Laufs 2014, Allen, Damaraju et al. 2017) have reported identification of potential electrophysiological correlates of fluctuations in BOLD FC thus suggesting the neurophysiological origin of FC and linkage to cognitive as well as vigilance states of the brain. Increasingly, information in the temporal variability of the correlation structure between RSNs is being leveraged to identify group differences between the diseased and healthy controls (Damaraju, Allen et al. 2014, Rashid, Damaraju et al. 2014, Miller, Yaesoubi et al. 2016). Analysis of the temporal dynamics of network time-courses, also referred to as dynamic functional network connectivity (dFNC), is generally carried out by applying a sliding window correlation (SWC), dynamic conditional correlation (DCC), phase synchronization (PS), co-activation patterns (CAPs) or a time-frequency coherence (TFC) approach. The SWC method (Sakoglu, Pearlson et al. 2010, Allen, Damaraju et al. 2012) evaluates temporal FC by calculating the correlation between the time-courses of the components of interest at all time-points within a chosen window, and repeating the process by gradually moving the window through the scan length. Recently introduced to the neuroimaging community, the DCC method (Lindquist, Xu et al. 2014, Choe, Nebel et al. 2017) is a multivariate volatility model that estimates model parameters through quasi-maximum likelihood methods and is widely used in the finance literature to estimate time-varying variances and correlations. The PS method (Glerean, Salmi et al. 2012) involves comparison of two signals by separating the amplitude and phase information parts, and has been reported to have maximal temporal resolution; however, its use is limited to narrow band signals. The focus of the CAPs method (Liu, Chang et al. 2013, Liu and Duyn 2013) is to identify and study instantaneously co-activating patterns. Lastly, the TFC approach is an extension of the coherence and time-domain approaches that features connectivity pattern estimation using the frequency and phase lag information, and has been successfully used in studying connectivity in a few brain regions of interest (Chang and Glover 2010) or whole-brain connectivity (Yaesoubi, Allen et al. 2015). In this paper, we will focus on using the sliding window method as it applies minimal assumptions and few data transformations. FC analysis using sliding window is often proceeded by a rigorous FC “state” profile estimation and characterization process (Allen, Damaraju et al. 2012), wherein the FC states are referred to the distinct discrete, transient patterns of FC, conceptually analogous to the quasi-stable EEG microstates. The estimated state profiles represent transient patterns of functional connectivity that vary over time; it was found in Allen, Damaraju et al. (2012) that subjects tend to remain in the same state profile for long periods of time before transitioning to one of the other state profiles (after multiple TRs).

The term “chronnectome” was recently introduced at the Mind Research Network to describe a focus on identifying dynamic i.e. time-varying, but reoccurring, patterns of coupling among brain regions. The chronnectome could be thought of as a brain model in which connectivity patterns as well as nodal activity varies in fundamental modes through time. The chronnectome project, aimed at standardizing methods evaluating dynamic FC with an ultimate goal of working towards evaluation of association of FC to physiologically meaningful changes, had hypothesized the existence of canonical patterns of dFNC associated with the resting state of the human brain. Multiple dFNC frameworks in the chronnectome project have evaluated dynamics of the FC SPs through different methods, but have commonly reported the FC SPs to be stably present in the data, highly structured, and reoccurring over time (Allen, Damaraju et al. 2012, Miller, Yaesoubi et al. 2016).

The need to examine the reliability of the emergent discrete FC patterns has been mentioned in several reviews on this emerging field (Hutchison, Womelsdorf et al. 2013, Calhoun, Miller et al. 2014, Calhoun and Adali 2016). Our initial work published as a short conference paper (Abrol, Chaze et al. 2016) evaluated replicability of the FC patterns emergent in the hard-clustering dFNC framework (Allen, Damaraju et al. 2012) over numerous large, age-matched and independent data samples from a huge dataset of 7500 human brain resting state fMRI scans. We found high levels of correlation in the SPs across these independent samples, which motivated us to further evaluate replicability in the dynamic FC. In the current paper, we test the replicability of the FC SPs by two frameworks: (1) building on our initial work in hard-clustering dFNC approach by additional testing for consistent clustering for a range of clusters, conducting surrogate dataset analysis to verify the driving factor of clustering, and examining additional separate-state summary measures such as fractional times and percentage occurrence time, and (2) extensively evaluating similarity in the derived across-state dynamic measures in the fuzzy meta-state dFNC framework (Miller, Yaesoubi et al. 2016), conducting an extensive battery of validation tests for this approach as well. The overall approach undertaken for this study is summarized below; detailed methodology can be found in the materials and methods section.

### 1.1. Summary of the Analytic Approach

Anomaly detection in the form of a correlation analysis on the five upper and lower slices of the preprocessed functional images was performed on all 7500 resting fMRI datasets in order to detect the scans that failed the reorientation process, or had any missing slices. To test our hypothesis, we pursued a data-driven approach (Figure 1) performing a high model order group-level spatial independent component analysis (gICA) on the subset of 7104 datasets (about 95% of the total data) that passed the above analysis, thus resulting in maximally spatially independent functional networks or RSNs corresponding to different anatomical and functional segmentations of the brain. Using a higher model order for spatial gICA enables us to extract multiple focal network nodes within multiple network domains (Allen, Erhardt et al. 2011, Allen, Damaraju et al. 2012) viz. sub-cortical, temporal, frontal, parietal, etc. The components emergent from this decomposition were used only as a spatial sorting template for the subsequent independent analyses. In the next phase, the first 7000 scans of the subset of 7104 datasets were partitioned into 28 age-matched independent samples of 250 subjects each, enabling us to evaluate replication on multiple independent datasets each with a relative large number of subjects. Each independent sample (referred hereon as group) was decomposed using the same gICA parameters, and subject time-courses and maps for all groups were subsequently back-reconstructed to perform functional connectivity analysis. Components across the multiple decompositions were mapped onto the template RSNs using a greedy spatial correlation analysis. The reconstructed time-courses went through additional processing to remove probable noise sources and outliers, and subsequently the processed time courses underwent dFNC analysis with a sliding window method thus resulting in emergence of inter-ICN covariance patterns for the different time windows. To explore structure and frequency of these estimated windowed connectivity patterns (CPs), a state estimation and characterization process was implemented using a hard clustering approach (Allen, Damaraju et al. 2012) as well as a fuzzy meta-state approach (Miller, Yaesoubi et al. 2016).

**Figure 1:**
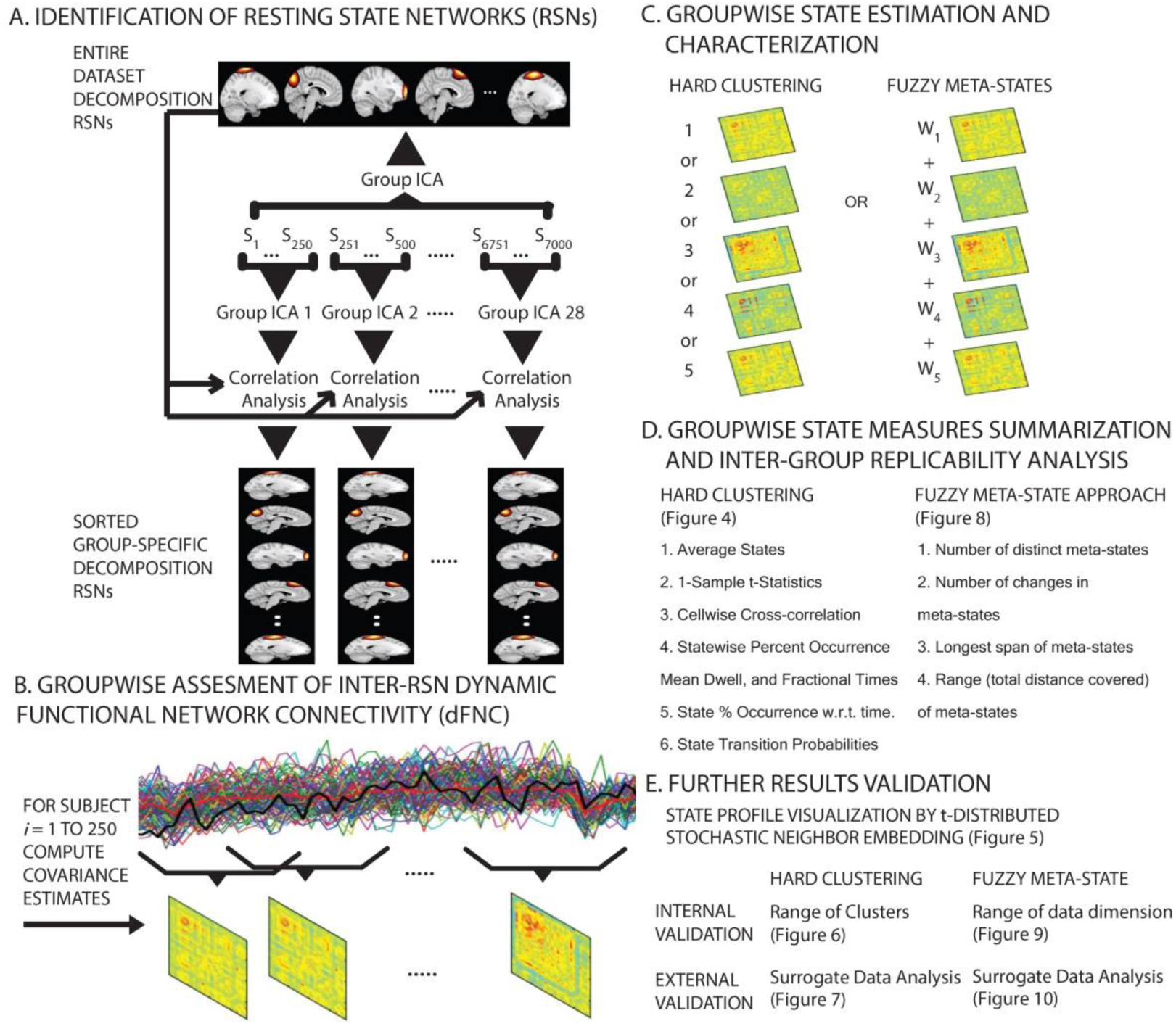
Summary of the research methods. (A) 61 components from the Group Independent Component Analysis (GICA) on the entire dataset were identified as reference Resting State Networks (RSNs). The first 7000 datasets were partitioned into 28 age matched, independent groups each having 250 scans. All partitioned samples underwent GICA using the same parameters as the entire dataset GICA. Spatial maps and time-courses of the individual subjects were reconstructed for all decompositions. Networks across the decompositions were mapped onto the reference RSNs using a greedy spatial correlation analysis, and 37 components over a fixed threshold were retained for dFNC analysis. (B) The reconstructed time-courses went through additional processing to remove any probable noise sources and outliers. These processed time courses underwent dFNC analysis with a sliding window method thus resulting in emergence of inter-RSN covariance patterns for the different time windows. (C) To explore structure and frequency of the windowed connectivity patterns (CPs), recurring connectivity state profiles (SPs) were estimated and characterized using a hard-clustering approach (CPs clustered to map one of the recurring connectivity SPs), and a fuzzy meta-state approach (CPs decomposed into meta-states). (D) Sorted SPs were summarized deriving several similarity statistics in both approaches to evaluate replicability. (E) Results were validated using extensive internal (using the original fMRI data) as well as external (using synthesized surrogate datasets) validation methods in both approaches.

We first applied the hard-clustering approach on the windowed CPs to identify the reoccurring basic SPs. This approach makes use of a clustering algorithm to assign the high dimensional CPs to one of the clusters, and has been successful in classifying patients from healthy controls (Rashid, Damaraju et al. 2014). However, this approach maps the high-dimensional CPs to one dimension i.e. they are allocated membership of one of the clusters. With existence of hard defined boundaries, distant CPs may still be assigned the same centroid, whereas lesser dynamically different CPs may be assigned two different clusters. Such an approach is complemented by a more flexible, fuzzy framework of expressing connectivity in the state estimation and characterization process. So, our second approach is based on a meta-state framework proposed recently to allow a subject’s state to be represented by varying degrees of multiple states, and is claimed to exhibit lesser distortion in the CPs and other features under investigation since contributions of all overlapping states are recorded (Miller, Yaesoubi et al. 2016).

For each independent group, state measures were summarized separately using k-means clustering for a range of clusters in the hard-clustering approach, whereas state measures were summarized across all states using multiple decomposition techniques including temporal ICA (tICA), spatial ICA (sICA) and principal component analysis (PCA) in addition to (fuzzy) k-means clustering in the meta-state approach. Furthermore, extensive group analysis was carried out on synthesized surrogate datasets to test whether the covariance structures in the RSN time-courses were indeed the actual cause of the state profiles and different from the covariance structures of the randomly shuffled RSN time-courses. Finally, we studied the impact of head motion in our analyses by making a comparison of the replicability metrics in observed dynamic measures with the replicability metrics from de-spiked and motion-regressed dynamic measures. Overall, results from this study strongly suggest faithful reflection of dFNC properties and reproducibility of the derived statistical measures across the independent partitions.

## 2. MATERIALS AND METHODS

### 2.1. fMRI Data Acquisition and Preprocessing

This study worked with resting state data that was previously collected, anonymized, and had informed consent received from subjects, both healthy and patients (aged between 13 to 75 years), as per the institutional guidelines practiced at the University of New Mexico (UNM) and the University of Colorado Boulder (UC, Boulder).

All 7500 resting state scans chosen for this analysis were acquired using 3-Tesla Siemens TIM Trio MRI scanners with 12 channel radio frequency coils at the Mind Research Network (MRN) in association with UNM, or using the same hardware scanner at UC, Boulder. Both scanners used the exact same acquisition parameters (except for the repetition time) for most of the subjects. T2^∗^-weighted functional images were acquired using a gradient-echo EPI sequence with TE = 29 ms, TR = 2s (6992 scans) or 1.3s (8 scans), flip angle = 75°, slice thickness = 3.5 mm, slice gap = 1.05 mm, field of view = 240 mm, matrix size = 64 × 64, voxel size = 3.75 mm × 3.75 mm × 4.55 mm. The sampling rates of the scans were matched before the dFNC analysis. The scans had variable length with the minimum scan length being 150 TRs; however, only the first 150 time-points of all scans were studied. This data was a de-identified convenience dataset for which we do not have access to the health and identifier information. While it would be useful to have that information and evaluate possible subgroups and additional variables of interest, our perspective was that having this additional variability should, if anything, make the possibility of replicating the state patterns even less likely.

The functional data were preprocessed using MRN’s automated preprocessing pipeline based on the SPM software. The data pre-processing pipeline integrated removal of the first three images in the scans to avert T1 equilibration effects, realignment using INRIalign, timing correction of slices with the middle slice fixed as reference, spatial normalization of data into the Montreal Neurological Institute (MNI) space, re-slicing of data into cubic voxels of side 3mm, and data smoothing using a Gaussian Kernel with the full-width at half-maximum (FWHM) set to 10 mm.

Anomaly detection in the form of a correlation analysis on the five upper and lower slices of the functional images was performed on all 7500 scans in order to detect scans that failed the reorientation process, or had any missing slices. This outlier detection removed 396 subjects, thus leaving behind a total number of 7104 subjects corresponding to approximately 95% of the available data.

### 2.2. Group ICA and Postprocessing

The built-in auto-masking function in the AFNI software was leveraged to create an average mask to be used as the template mask input while running group ICA on the data that passed the anomaly detection analysis. In a multistep procedure to identify the RSNs, data was decomposed into maximally spatially independent components using functions from the Group ICA of fMRI Toolbox (GIFT). With a higher model of 100 (aiming at finer parcellation), this group decomposition used only the initial 150 time-points of all scans.

Choosing a higher number of principal components at the subject level stabilizes back-reconstruction and retains maximum variance in the data as shown in (Erhardt, Rachakonda et al. 2011). So, in the group ICA analysis, the entire dataset was transformed into 130 principal components using standard principal component analysis (PCA) at the subject level in the first data reduction step retaining maximum subject-level variance (greater than 99.99%), and further down to 100 components by implementing group level PCA in the second data reduction step. ICASSO (Himberg, Hyvarinen et al. 2004) was used to investigate reliability of the estimated independent components, and it was found that the estimates exhibited tight clustering, hence converging consistently amongst several runs. The spatial maps and time-courses of the individual subjects did not undergo backward reconstruction since it was not required for this specific analysis. Careful analysis on the emergent decomposition patterns confirmed 61 components having no correspondence to any known imaging, physiological, movement-related artifacts. Component map templates for these shortlisted components were assessed and distributed into the somatomotor, parietal, frontal, default mode, visual, temporal and cerebellar networks (Figure 2).

**Figure 2:**
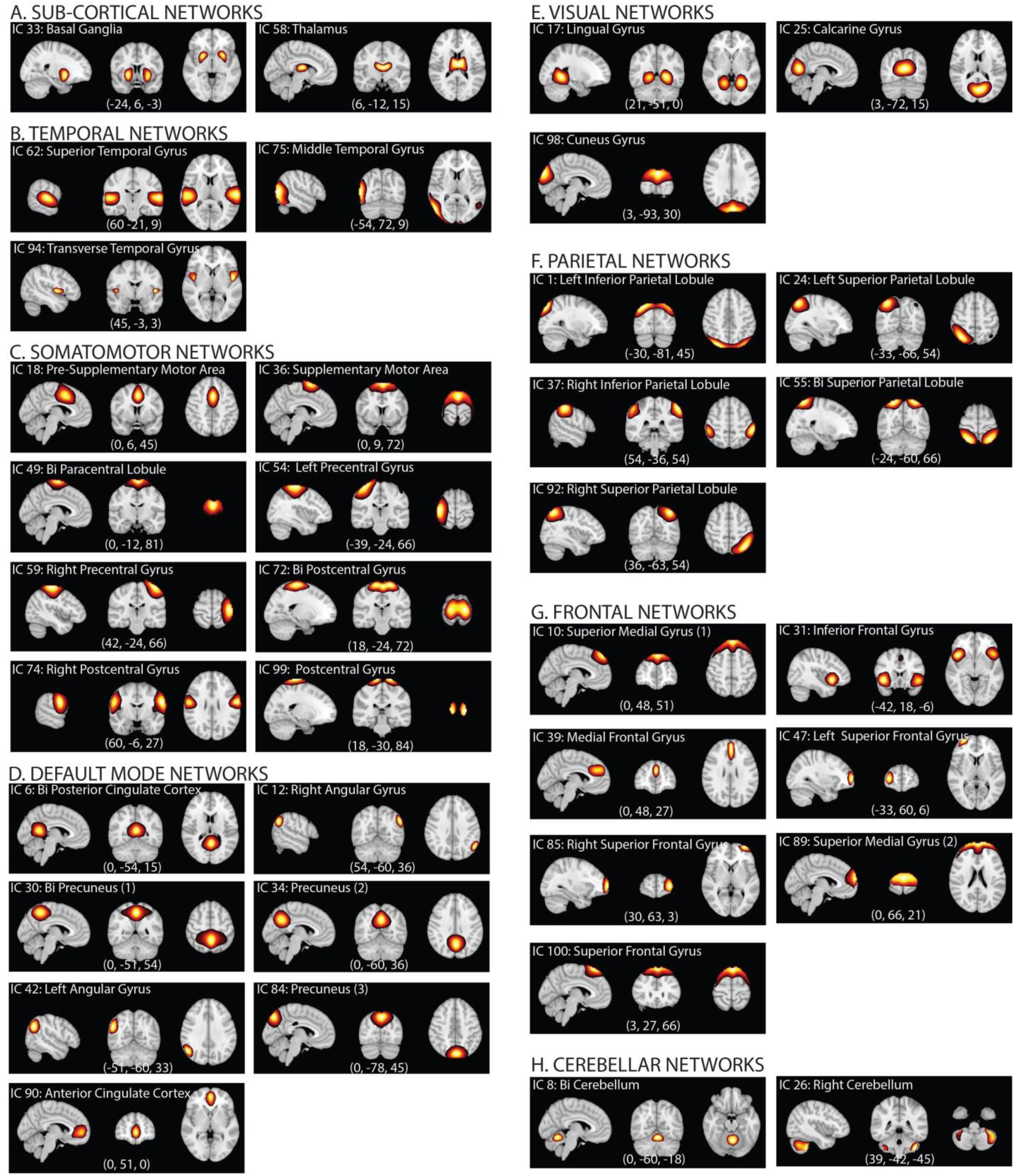
Resting State Networks (RSNs). Spatial maps of the 37 retained RSNs at the most activated sagittal, coronal and axial slices.

In an effort to set up the maximum possible number of independent samples each having a large partition size, the first 7000 of the 7104 scans were partitioned into 28 age matched groups each having 250 scans. These age-matched groups were a mix of subjects from both sites (UNM: a total of 6472 subjects, with 231.1 +−4.3 subjects per group, all subjects with a TR of 2 seconds; UC: a total of 528 subjects with 18.9 +−4.2 subjects per group, 520 subjects with a TR of 2 seconds), and had an average age of 31.65 years with an average standard deviation of 13.8 years for subjects within the groups.

With a focus on evaluating repeatability of the dFNC metrics corresponding to the partitioned samples, all the samples underwent separate group ICA decompositions. Similar to the entire dataset group ICA decomposition, standard PCA was performed at subject level for reducing data down to 130 components in the first step, and further down to 100 components by using group level PCA in the second step. Again, alike the entire dataset decomposition, ICASSO (Himberg, Hyvarinen et al. 2004) was used to verify consistency of the estimated independent components in all 28 group decompositions. However, subject specific time courses and spatial maps were also back reconstructed for these group decompositions since they were required for the inter-component correlation analysis.

The reconstructed component time-courses went through additional processing steps to remove any residual noise sources mostly including low frequency trends originating from the scanner drift, motion related variance emerging from spatial non-stationarity caused by movement, and other non-specific noise artifacts unsatisfactorily decomposed by the implemented linear mixed model. More specifically, the post-processing steps featured de-trending existing linear, quadratic and cubic trends, multiple linear regression of all realignment parameters together with their temporal derivatives, outlier detection using 3D spike removal, and low pass filtering with high-frequency cut-off being set to 0.15 Hz. Lastly, the time courses were variance normalized which meant the covariance structures from the sliding window approach were equivalent to the correlation structures.

### 2.3. Component Selection

An extensive evaluation of the spatial maps and spectral composition of the components resulting from the entire dataset gICA decomposition was carried on to identify physiological non-artifactual, previously established networks. Specifically, 61 template components with local peak activations in gray matter, time-courses dominated by low-frequency fluctuations, and high spatial overlap with known RSNs were selected for further analysis. For each of the 28 group decompositions, respective components were mapped to the identified 61 non-artifact template components from the entire dataset decomposition by finding best unique matches through a greedy correlation analysis. The 37 components with highest correlation values or more specifically above the first quartile correlation threshold value of 0.65 and global correlation threshold value of 0.4 for all sample decompositions were retained for the dynamic FNC analysis.

### 2.4. FC Estimation and temporal Variability

Dynamic FNCs between all 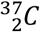 (666) RSN pairs in each of the 28 group decompositions were estimated using a tapered sliding window featuring convolution of a rectangular window (width = 30 TRs = 60 seconds) with a Gaussian (σ = 3 TRs), and subsequently sliding this tapered window in gradual steps of 1 TR, finally resulting in as many as W = 120 windows. Hence, for each group, dFNC was estimated subject wise to get a series of 120 correlation vectors corresponding to the series of windowed partitions of the subject specific time-courses.

#### 2.4.1. Approach 1: Hard Clustering

In this approach, the frequency and structure of the reoccurring 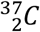 (666) dimensional dynamic windowed CPs emerging from all subjects in a specific group was modularized by implementation of the classical hard k-means clustering. The clustering algorithm was implemented using the Manhattan (cityblock) distance as the similarity measure since the L1 norm has been suggested to be a more effectual similarity measure than the L2 norm for high dimensional data.

The elbow criterion was used to derive the number of clusters input to the clustering algorithm. In this method, the central idea is to run k-means for different values of a specified number of clusters (k), and determine the case that maximizes within-cluster similarity and between-cluster dissimilarity concurrently. More specifically, we measured the ratio of within cluster sum of squared distances (dispersion in the cluster) to the sum of squared distances for all other observations (total variability outside that cluster). Finally, we evaluated this measure averaged over all clusters with respect to the number of clusters, and validated the case after which the gain in explanation of variation in data made only a marginal difference.

Furthermore, a two-level clustering was implemented in an effort to reduce the clustering error where an initial point input to the second level clustering was estimated in the first level clustering, and all windowed FNC data was clustered in the second level clustering. The initial point input was found by estimating and clustering the subject exemplars (corresponding to subject FNC windows featuring highest variance in FNC). More specifically, for each subject time-point (window), the standard deviation in FNC was computed, and windows at the time-points exhibiting local maxima were retained as subject exemplars and subsequently clustered. The centroid connectivity patterns resultant from this first level clustering were then set as the initial point input to the second level clustering of all FNC data. This two-level clustering process is similar to EEG microstate analysis (Pascual-Marqui, Michel et al. 1995), and was thoroughly tested for consistency for fMRI data in Allen, Damaraju et al. (2012). Repeating the initial as well as final clustering 150 times to increase the likelihood of escaping local minima, stable connectivity SPs were obtained for each of the groups.

Connectivity SPs emergent from the group-wise clustering analysis contain information on inter-RSN connectivity strength and variation in a particular group, and can be thought of as states that the subjects repeatedly transit into through the course of the scan. To evaluate replicability of state measures across groups, all sets of SPs were first sorted across groups in a multiple step greedy similarity analysis using Manhattan distance as the similarity measure. In each step, a new group was fixed as a reference to which the remaining groups where evaluated for similarity and then sorted according to the similarity distance thus eventually resulting in 28 sets of sorting orders. In the final step, the statistical mode over this structure of best matches of SPs was validated as the final sorting order. The least frequency of any of the modes was observed to be 22 out of the 28 groups, and similar results were achieved by using other L1 and L2 (Euclidean, squared Euclidean, correlation distance) similarity measures in the clustering algorithm, thus confirming reliability in the sorting process. Summary measures as discussed in the results sections were computed and compared across the sorted SPs in this clustering approach.

Visualizing data is considered important for quality control in any field, and hence we made an attempt to visualize the projections of the high dimensional CPs onto a two dimensional space by using the tSNE algorithm (Maaten and Hinton 2008). In the tSNE projection analysis, Euclidean distance between points is computed, and modelled as conditional probabilities with which one point would pick another as its neighbor such that more similar points are located nearby. Data is preprocessed with PCA reducing dimensionality to initial number of dimensions at the start of the learning. Perplexity of Gaussian distributions in higher dimensional space can be interpreted as the smoothing measure of number of effective neighbors. In this projection analysis, the initial number of dimensions = 50, initial learning rate = 500, number of iterations = 1000, and Gaussian perplexity was set to 50.

#### 2.4.2. Approach 2: Fuzzy Meta-states.

Computation of meta-states involves derivation of the windowed connectivity correlation data in a similar fashion as in the hard-clustering approach. In this approach, the 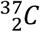 (666) dimensional windowed FNC covariance structures were decomposed into fewer dimensional (o) connectivity patterns (CPs) using one of the commonly used data-driven approaches viz. temporal ICA, spatial ICA, k-means and PCA. The lower dimensional CPs are maximally mutually independent time-courses with overlapping connectivity profiles in case of temporal ICA decomposition, maximally independent spatial patterns in case of spatial ICA decomposition, and orthogonal projections capturing maximal variance for the PCA decomposition. In case of the k-means clustering approach, cluster memberships are assigned to get low within cluster distances and high between cluster distances with the cluster centroids being treated as basis correlation patterns.

The windowed data decomposition was followed by assessment of contributions of the emergent, maximally independent patterns to the actual windowed correlation CPs. Finally, the real-valued weights associated with these states were estimated for every windowed FNC pattern, and the discretized version of this lower dimensional (o = 5 for main discussion, and o = 2 through o = 5 for comparison of results) characterization of the 666-dimensional CPs was achieved with a signed quartile transformation which resulted in meta-states. In our work, we compare results from the different decomposition methods, but mainly focus on the temporal ICA decomposition throughout the meta-state analysis discussion. The overall objective in this approach is again to calculate and compare group wise statistics from the meta-state profiles derived from all time windows of all the subjects in a given group.

## 3. RESULTS

In this section, we first describe results from the feature (or component) selection process following the group level ICA decompositions, and subsequently discuss findings from both dFNC approaches used in this study.

### 3.1. Feature Selection

The spatial maps of the 37 retained RSNs were thresholded (*t_c_*) by using mean (*μ_c_*) and standard deviation (*σ_c_*) parameters estimated using a normal-gamma-gamma (NGG) model (*t_c_* > *μ_c_* + 8*σ_c_*) to show regions contributing to the networks (Allen, Erhardt et al. 2011). The thresholded spatial maps of the RSNs at the most activated sagittal, coronal and axial slices are plotted in Figure 2. Co-ordinates of peak activations in MNI space, maximum activation level, activation region voxel volume, and associated Brodmann areas for the retained 37 RSNs are summarized in Supplementary Information (SI) Table 1. It can be easily confirmed that the retained RSNs demonstrate high similarity to RSNs from previous high-order decomposition studies (Allen, Erhardt et al. 2011, Allen, Damaraju et al. 2012).

### 3.2. Approach 1: Hard clustering

#### 3.2.1. Optimal clustering analysis

The elbow criterion used to derive the optimal number of clusters suggested an optimal number of five clusters for all groups (Figure 3a). For all groups and each k, the method was repeated 10 times as a consistency check. The group-wise boxplots of the validated number of clusters over the different runs are shown in Figure 3b.

**Figure 3:**
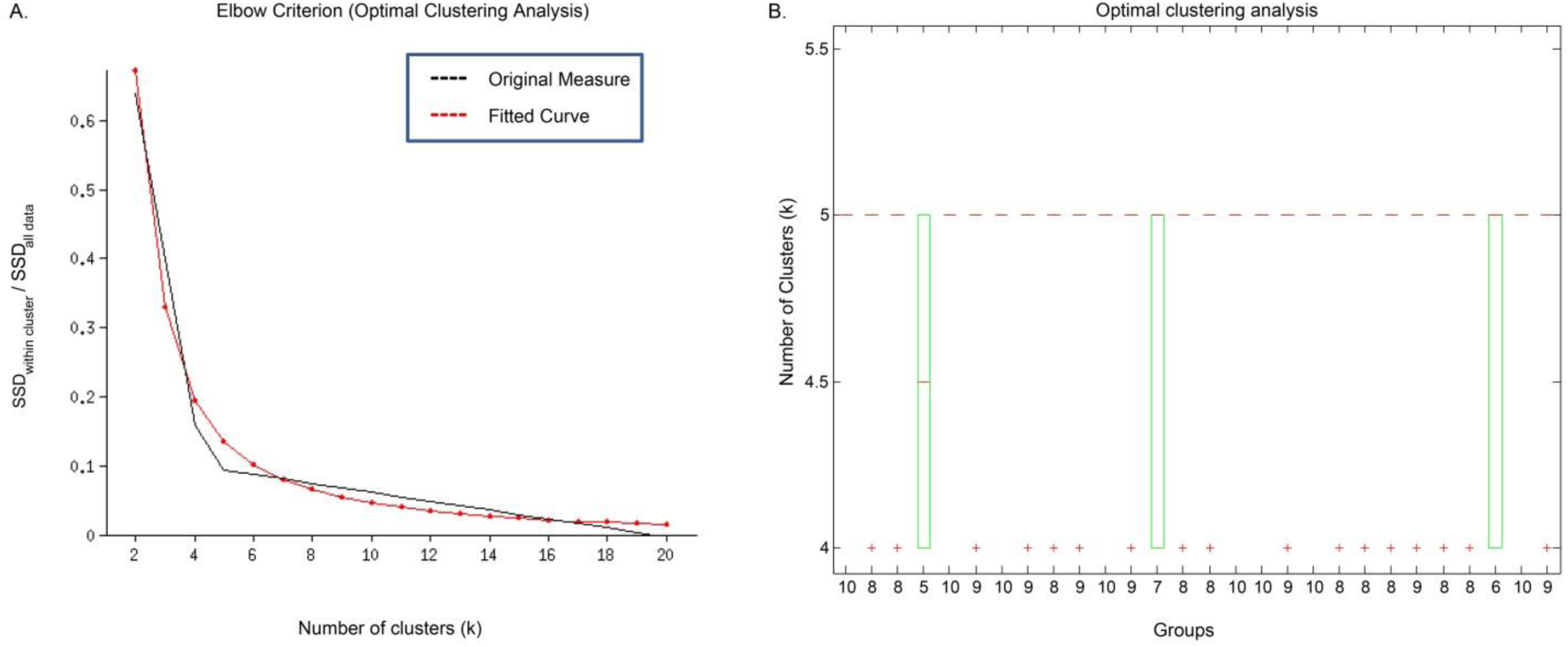
Optimal Clustering Analysis. (A) Elbow plot for a sample run; (B). Group-wise boxplots of the estimated optimal number of clusters over 10 independent runs. (B) The x-axis labels in Figure 3b illustrate the number of runs for that particular group (out of a total of 10 runs) that estimated the optimal value of k equal to 5. In all, 241 out of the 280 independent runs estimated the optimal value of k equal to 5; hence, this value of k was validated as the optimal clustering case for the rest of the study.

#### 3.2.2. State summary measures

Characterizing states and summarizing state metrics provides important and useful information e.g. the time spent in a particular state, the directional probability of transitions between two particular states, etc. that help us quantify replicability between the independent group decompositions. The state summary measures and similarity statistics evaluated in the hard-clustering approach are shown in Figure 4, and their relevance and contribution in eventually investigating replicability of the temporal dynamics is discussed subsequently.

**Figure 4:**
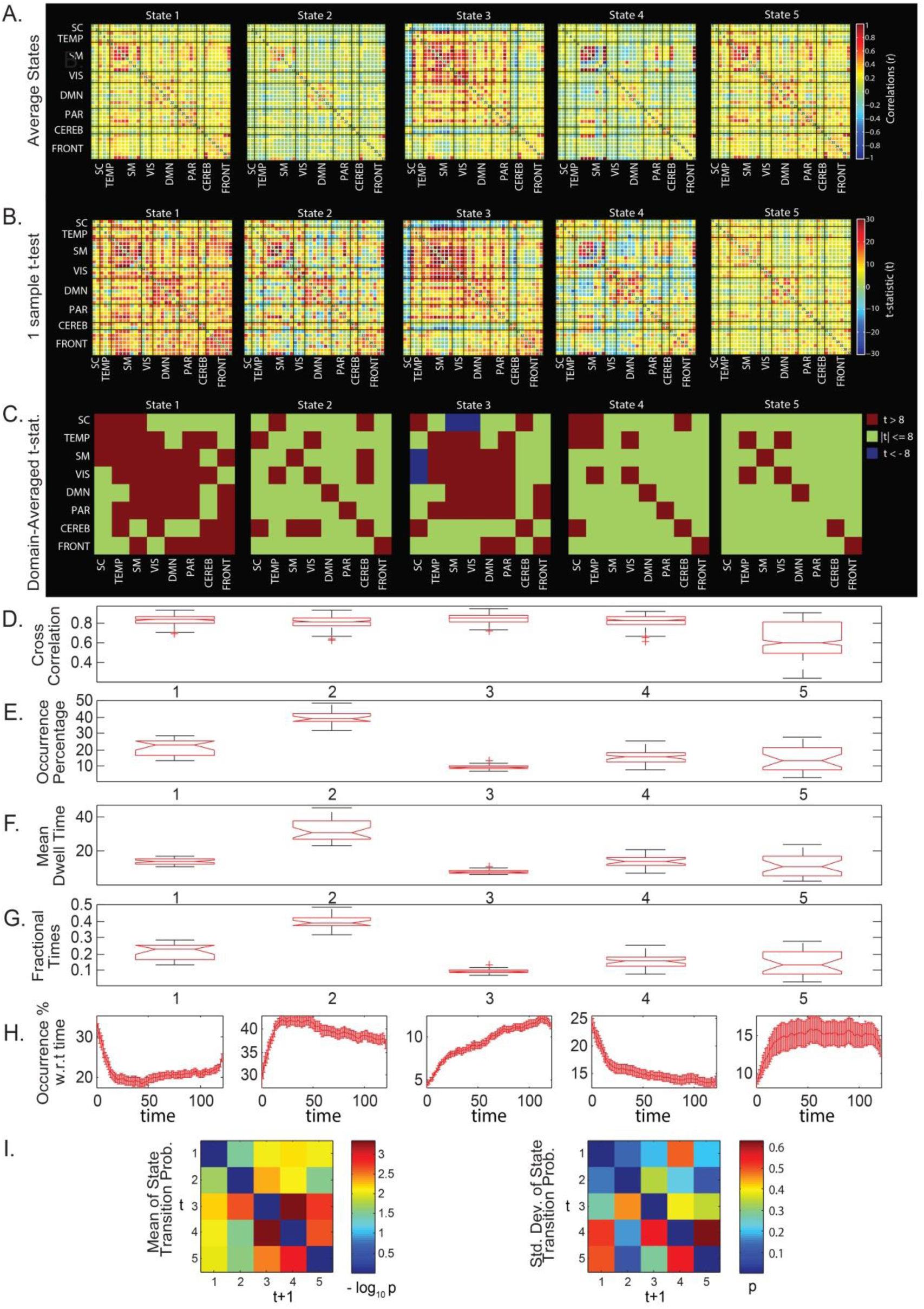
State summary measures in the clustering approach. (A) SPs averaged over all groups; (B) 1-sample t-test results on the SPs; (C) 1-sample t-test results averaged over the domains; (D) Boxplots of pairwise linear correlations of the SPs; (E) Boxplots of average occurrence % of the SPs; (F) Boxplots of the average mean dwell times of the SPs; (G) Boxplots of average fractional times of the SPs; (H) Occurrence percentages of the SPs modeled w.r.t. time; and (I) Mean and std. deviation of average state transition probabilities (modelled as a first order Markov chain).

The average state metric (Figure 4A) provides information on averaged connectivity and the percentage of occurrence considering all independent samples as one large sample, whereas the one-sample t-statistic metric (Figure 4B) highlights regions with high mean and smaller standard deviations, and hence the connections in the region can be considered to be more reliable. The 1-sample t-test statistics averaged over pairs of network domains can be seen in Figure 4C to highlight the most reliable network domain pairs in a particular state. Figure 4D shows the pairwise linear correlations of the mapped SPs across all the groups. Evidently, states 1 to 4 have high correlation numbers (first quartiles greater than 0.8) suggesting these states are highly reproducible across the independent samples, whereas state 5 with higher spread is not as fully reproducible as the other states. The considerably larger spread of state 5 in the correlation boxplot is explained in the t-distributed Stochastic Neighbor Embedding (tSNE) projection analysis (Maaten and Hinton 2008) in the coming sub-section where state 5 is actually observed as mixture of states 1 and 2. We also compare the state occurrence percentage, mean dwell time spent in each of the states, and fractional times for each of the states for all groups as shown in the boxplots in Figure 4E, 4F and 4G. For each of the boxplots, the group-wise state measures are well concentrated within their respective ranges with state 2 consistently observed as the most frequent (39.5 % average occurrence time), and hence with higher dwell and fractional times. The number of occurrences of the states is next modeled as a function of time (Figure 4H) so as to observe how the state occurrence frequencies increase or decrease with time. Due to the unconstrained nature of resting state, it is unlikely to obtain consistent temporal trends in the cognitive states of the brain. However, we could investigate existence of any consistent temporal trends in occurrence of the FC state profiles to motivate theories on their relation to vigilance, sleep or arousal states. Similar to earlier work (Allen, Damaraju et al. 2012), we observe a state with increased thalamocortical anti-correlation probably related to drowsiness (State 3) for which frequency of occurrence increases with time spent in the scanner and that occurs about 10% of the time in all groups. This observation is consistent with Tagliazucchi and Laufs (2014) who report reliable drifts between wakefulness and sleep during typical waking rest fMRI scans. Furthermore, EEG correlates suggest that this state corresponds to increase in low frequency delta and theta power suggestive of reduced vigilance (Allen, Damaraju et al. 2017).

Finally, state transition behavior is captured by a first order Markov model which helps in understanding the propagation of probability transitions through the network i.e. probabilities associated with entering or exiting a given state. For readability of this model, the average values of probabilities (*p*) of group-averaged state transitions across all groups have been transformed through a –*log*_10_*p* transformation. Hence, smaller values in the averaged state transition matrix (on the left in Figure 4I) correspond to high probabilities of transition from one state to another. The standard deviations in the averaged state transition matrices across all groups are shown to the right in Figure 4I. It can be observed that there is a high probability of being in the same state at the next time instant (high probability along the diagonals), as well as a high reliability of transition to and from state 2 as compared to other states as they tend to have higher mean transition probabilities and smaller standard deviations. However, transition probabilities, with high standard deviations in some cases, can also be highly variable.

#### 3.2.3. Visualizing state profiles

The tSNE algorithm is known to preserve the local structure of the data by projecting similar higher dimensional structures (with smaller pointwise distance) closer in the 2D space than the relatively distinct ones as the algorithm learns at a predefined learning rate while the data is being processed over a predefined number of iterations. The exemplar high-dimensional windowed FNC data from all of the 28 groups were projected onto a two-dimensional space using tSNE. The final projection of the exemplar data can be visualized in Figure 5A which suggests states 1, 2, 3 and 4 to be clustering consistently, whereas state 5 showed high variance and appeared more similar to states 1 and 2. This observation suggests that the 5^th^ state is not fully reproducible as the other states. Figure 5B shows the data for states 1 to 4 only (for all groups), and these 4 classes can be seen to be clustered in distinct, but touching regions. An assessment of the class conditional density peaks for these states in 2 dimensions (Figure 5C) and 3 dimensions (Figure 5D) revealed distinct density peaks for all 4 classes thus further supporting the existence of structure in the clustered data. Replicability of the states can be visualized from a different angle in SI Video 1 in which CPs from all data windows are projected a group at a time on top of data from groups already projected.

**Figure 5:**
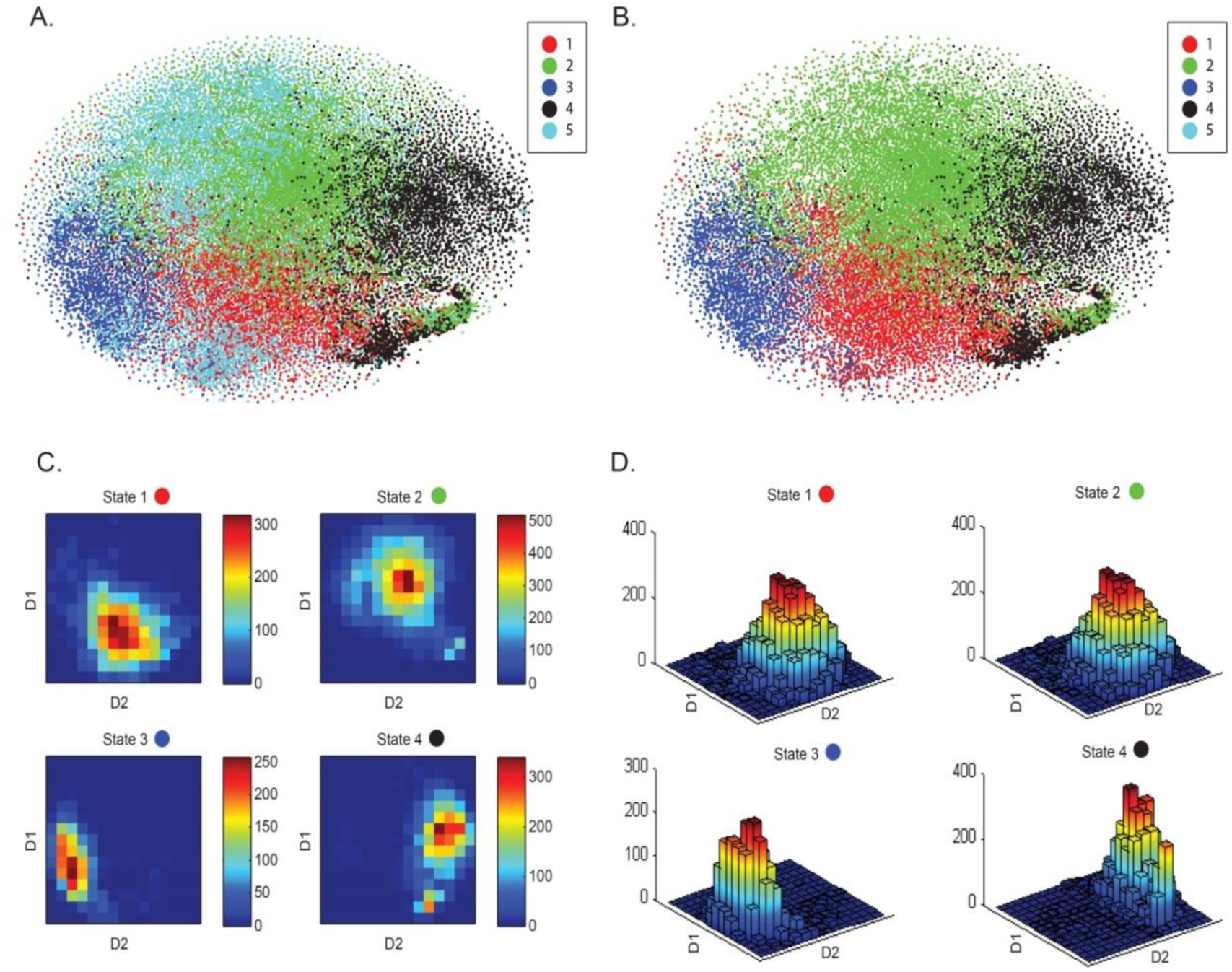
High-dimensional windowed FNC data projection onto a two-dimensional space using the t-distributed Stochastic Neighbor Embedding (tSNE) framework. (A) tSNE visualization of the windowed FNC data from all 28 groups suggests consistent clustering for states 1, 2, 3 and 4 for all groups, touching class boundaries and low degree of homogeneity for state 5. Additionally, from the state summary measures, State 5 was seen to be less reproducible as compared to the other 4 states. (B) tSNE visualization for states 1 to 4 for all groups confirms distinct (but touching) clustering regions for these different data classes. (C) and (D) Class conditional densities for the states 1 to 4 in 2 dimensions (Figure 5C) and 3 dimensions (Figure 5D) reveal distinct peaks for all 4 classes thus validating the structure in the data.

#### 3.2.4. Internal Validation: Clustering for a range of k

With an objective of using the same dataset to internally validate results from the hard-clustering approach, additional analysis was carried out for a range of number of clusters (k = 2 to 10). Figure 6 plots the state profiles over this range for the first group, whereas results for all the groups are presented in SI Video 2. It must be noted that within every group, the state connectivity profiles had to be sorted since the order of the centroids or clusters emergent from the k-means algorithm is not unique. After sorting through a greedy algorithm as defined in the methods section, it can be clearly seen that clustering results for different clustering indices are consistent.

**Figure 6:**
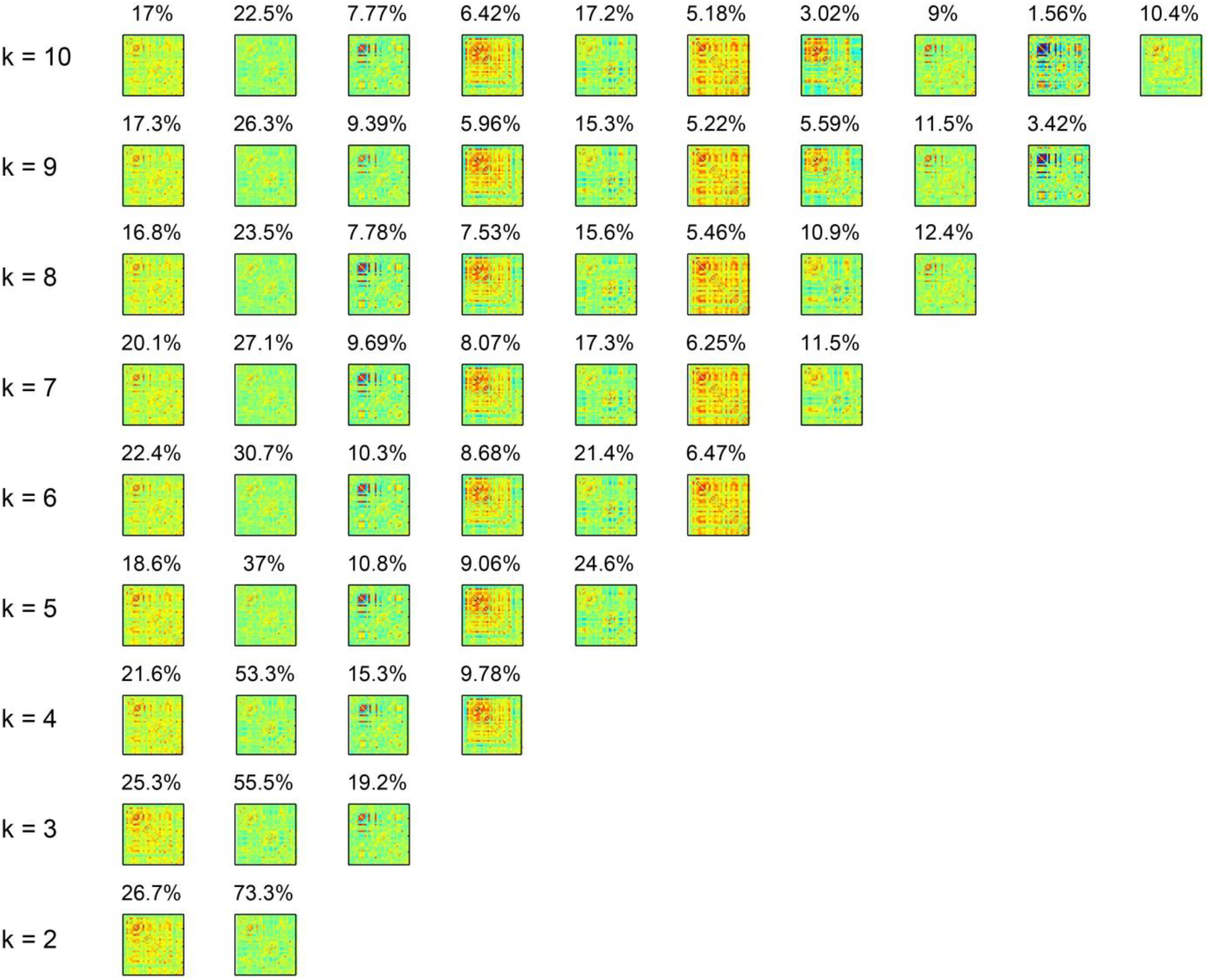
Internal validation in the hard-clustering approach. Clustering results for a range of number of clusters (k = 2 to 10) demonstrate high similarity of the emergent state profiles.

#### 3.2.5. External Validation: Clustering surrogate data

To verify the driving factor of the emergent discrete FC state profiles (SPs), we explored surrogate data testing by phase randomization (Prichard and Theiler 1994) of the RSN time-courses. Similar to the phase randomization procedure used in Damaraju, Allen et al. (2014), Handwerker, Roopchansingh et al. (2012) and Hindriks, Adhikari et al. (2016), the surrogate RSN time-courses were generated by Fourier transforming the RSN time-courses estimated from real fMRI data, adding a uniformly distributed random phase to each frequency in this frequency domain data, and finally inverse Fourier transforming the frequency domain data back to the time-domain. Adding the same random phase to the same frequency components of the RSNs preserves the static FNC and the lagged cross-covariance structure in the surrogates. This class of surrogates, hereinafter referred to as “consistent” phase randomized (CPR) surrogates, correspond to the null hypothesis that the real RSN time-courses are explained by a linear, stationary Gaussian process (Schreiber and Schmitz 2000, Borgnat, Flandrin et al. 2010, Richard, Ferrari et al. 2010, Liegeois, Laumann et al. 2017). Alternatively, adding different random phases to the same frequency components of the RSNs disrupts the static FNC and the lagged cross-covariance structure in the surrogates. This alternate class of surrogates, hereinafter referred to as “inconsistent” phase randomized (IPR) surrogates, instead correspond to the null hypothesis that the real RSN time-courses are explained by a linear Gaussian process with static FNC approximately equal to zero (Hindriks, Adhikari et al. 2016). In this initial part of the analysis, we will be using the IPR surrogates only to seek an explanation to clustering as the null generated from this class is not appropriate to make inferences about stationarity of the observed data since lesser than required properties of the observed data are preserved by construction. It must also be noted that by construction of surrogate RSN time-courses, the mean, variance and power spectrum of both surrogate classes are identical to that of the real RSN time-courses, and subsequently by the Weiner-Khintchine theorem, both surrogate classes will have the same temporal autocorrelation as the real RSN time-courses.

100 CPR and 100 IPR surrogate datasets for the real RSN time-courses were generated each of which underwent dFNC analysis and subsequent clustering individually. The SPs emergent from the different surrogate datasets were mapped to the SPs in the real fMRI dataset, and finally a scalar correlation distance (averaged across the SPs) was computed for each surrogate dataset. Figure 7A illustrates the distributions corresponding to the two surrogate classes where it can be seen that the CPR surrogates show a very small correlation distance (very high correlation), and the IPR surrogates show very large correlation distance (very low correlation) as compared to the real data SPs. This observation suggests that clustering is substantially driven by the lagged cross-covariance structure of the RSN time-courses and not solely by the linear autocorrelation structure of the RSN time-courses or dissimilarities in mean and variance across subjects.

**Figure 7:**
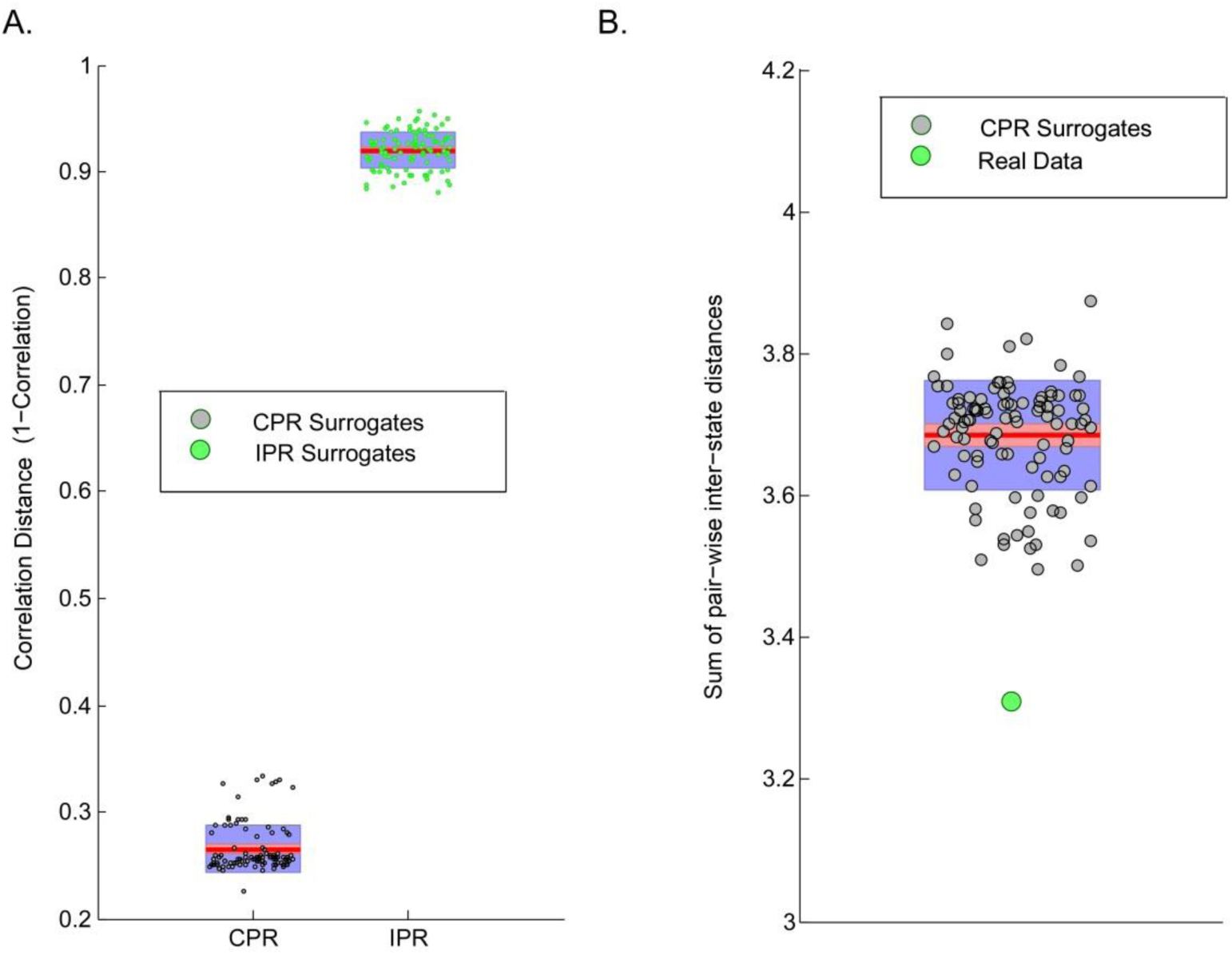
External Validation in the hard-clustering approach. (A). SPs emergent from clustering windowed FNC data corresponding to real fMRI data exhibit high correlation with SPs from similar analysis on 100 synthesized surrogate datasets of RSN time-courses with consistent phase randomization (CPR) and low correlation in case of inconsistent phase randomization (IPR); and (B) Observed sum of pair-wise inter-state distances in real data in comparison to the null distribution of this test statistic approximated from 100 CPR surrogate datasets.

Additionally, the presence of any significant differences in the statistical measures from the real and CPR surrogate data was explored by approximating the null distribution for a test statistic, namely sum of pair-wise inter-state distances, from the multiple CPR datasets, and subsequently comparing the observed value of this statistic for real data against the generated null. The CPR null was rejected for this test statistic in all groups which suggests presence of non-Gaussianity, non-linearity or non-stationarity, or a combination of these properties in the observed time-courses. Unfortunately, further non-trivial testing is required to narrow down on the cause of the rejection of the CPR null, a topic out of scope of the focus of this study and worth exploring in the future. Nonetheless, the two results in Figure 7 jointly suggest that clustering was substantially, but not completely, explained by the lagged cross-covariance structure of the RSN time-courses.

### 3.3. Approach 2: Fuzzy meta-states

#### 3.3.1. State summary measures

To evaluate replicability in the group statistics, several meta-state metrics such as number of distinct meta-states occupied (n), number of switches in meta states (s), longest state span (r: largest L1 distance possible in occupied meta-state vectors) and finally the total distance covered by a subject (d: sum of L1 distances covered by a subject) are computed from the emergent meta-states. The group-wise histograms of subject meta-state metrics from the temporal ICA decomposition method as plotted in Figure 8A show similar spread and distribution across the groups. Figure 8B, the mean stem plots and standard deviation boxplots suggest low variation in the estimated group summary metrics (*σ_S_* = 1.0015, *σ_n_* = 1.0678, *σ_r_* = 0.5373, *σ_d_* = 4.5790). Similar results from alternative decompositions such as spatial ICA, PCA, and k-means clustering (Figure 8C) confirm the low variation observed in temporal ICA decomposition metrics. It must be noted that k-means clustering uses only 4 discrete states (1 to 4), and hence has dissimilar numbers as compared to the other three decompositions with a maximum 8 possible states (−4 to −1, 1 to 4). Nonetheless, meta-state metrics from the k-means decomposition are similar across the different groups showing low variation for the estimated dynamic measures (*σ_S_* = 1.3192, *σ_n_* = 1.0840, *σ_r_* = 0.5033, *σ_d_* = 1.6242).

**Figure 8:**
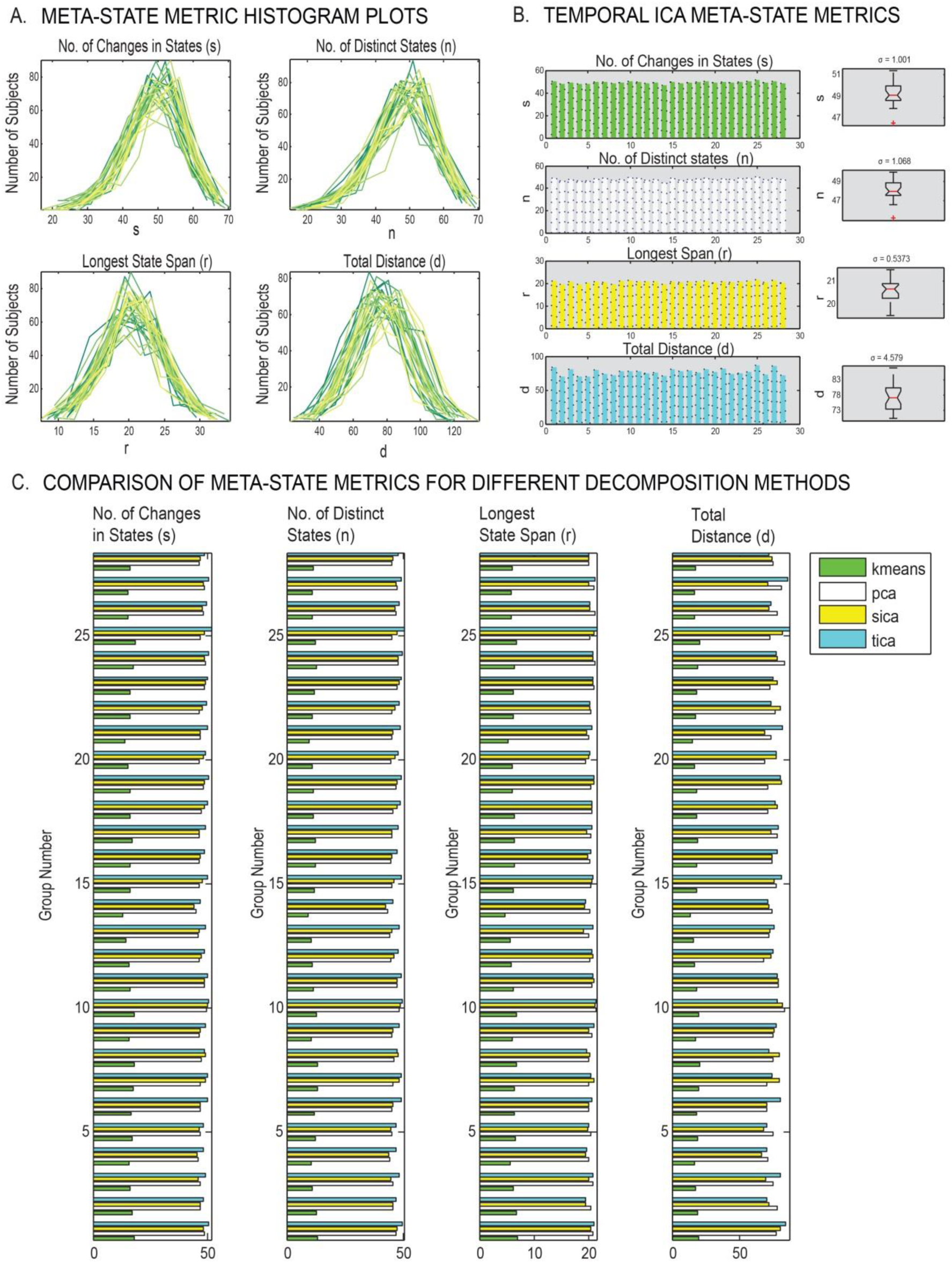
State summary measures in the meta-state approach. (A) Histogram plots of the estimated subject-specific temporal ICA meta-state metrics demonstrate similar distributions across all groups; (B) Boxplots of group-wise averages of temporal ICA meta-state metrics indicate low variation in group summary metrics; (C) Similarity of group summary metrics across different groups within and across different decomposition methods. Notably, the metrics are consistent across groups in k-means, but different from other methods since k-means uses only 4 discrete states (1 to 4), as compared to other methods that use 8 states (−4 to −1 and 1 to 4).

#### 3.3.2. Internal Validation: Testing range of dimensions

Sensitivity of the replicability results to the number of dimensions (model order o) in the meta-state approach is tested by performing the meta-state analysis with this parameter ranging from 2 to 5. As evident from the bar graphs and low standard deviations in Figure 9, there is great similarity in all meta-state summary metrics in all model orders, which further substantiates evidence of the replicability of the meta-state approach summary metrics across the independent samples. Notably, the averaged meta-state statistics for the entire dataset increase with the number of dimensions since the range of possible meta-states in the state space increases with the model order; however, within a given model order, high group-wise similarity in the dynamic measures is observed.

**Figure 9:**
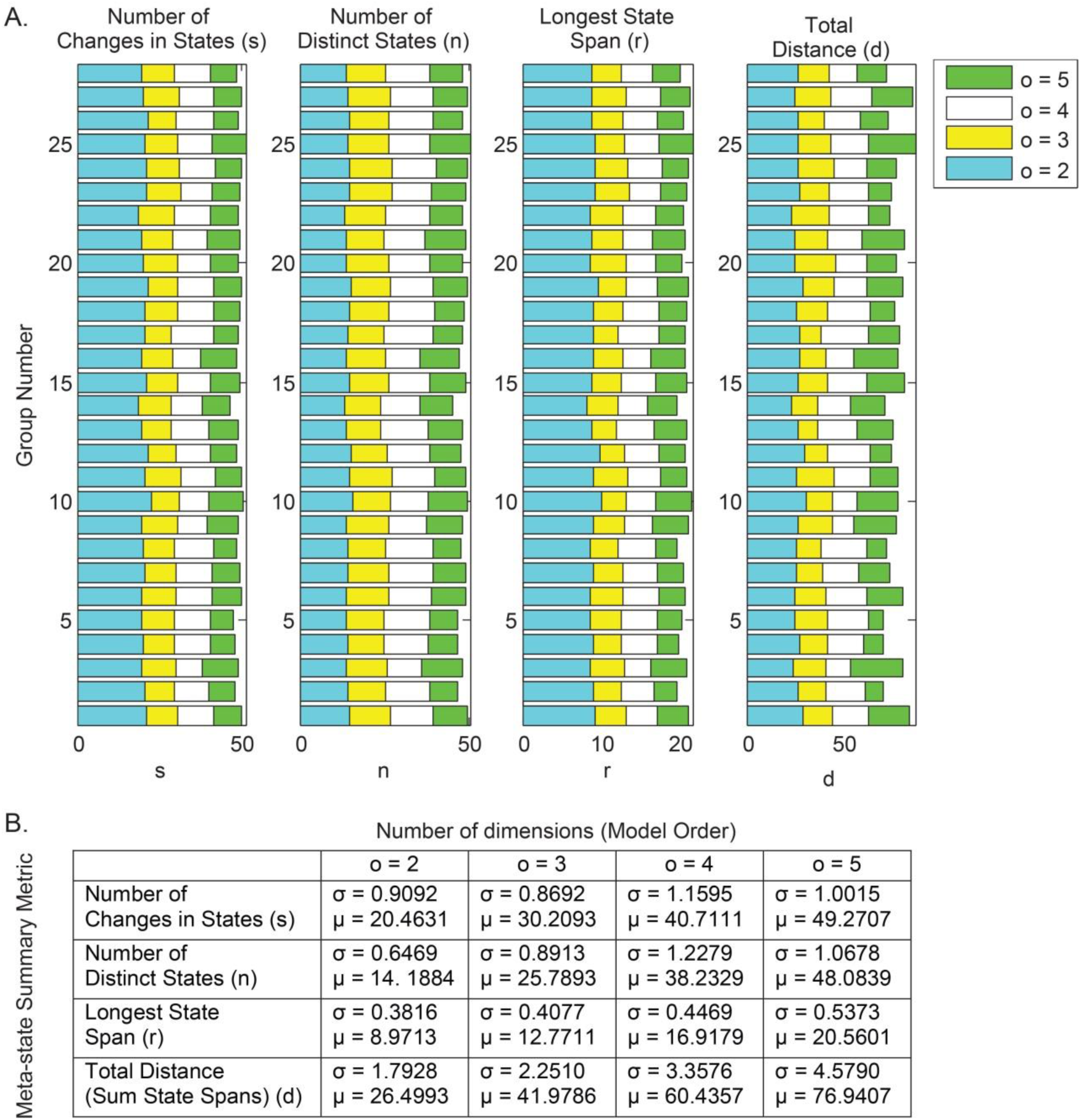
Internal Validation in the meta-states approach. (A) Sensitivity test of number of dimensions (o = 2 to 5) to the meta-states framework validates similarity in group summary measures; (B) Averaged metrics are consistent across groups, but increase with model order as range of meta-states is proportional to model order.

#### 3.3.3. External Validation: Decomposing surrogate data

Correspondence of the meta-state summary metrics to the RSN time-courses corresponding to real fMRI data was tested by meta-state permutation testing on a set of 100 CPR surrogate datasets of RSN time-courses. For all meta-state metrics, the outcome from the real dataset was determined and compared against the null distribution for the respective meta-state metrics generated from the different surrogate datasets. Figure 10 shows that all summary metrics for the real dataset are located completely outside the synthesized null distribution for the temporal ICA, spatial ICA and PCA decomposition methods. Similar results were observed for the k-means decomposition method as well. This rejection of the CPR null model for the studied meta-state metrics for different decomposition methods adds further evidence to presence of non-Gaussianity or non-linearity or non-stationarity, or any combination of these three properties in the observed RSN time-courses. Spotting the specific cause of rejection of the CPR null hereon is non-trivial and an interesting topic for future.

**Figure 10:**
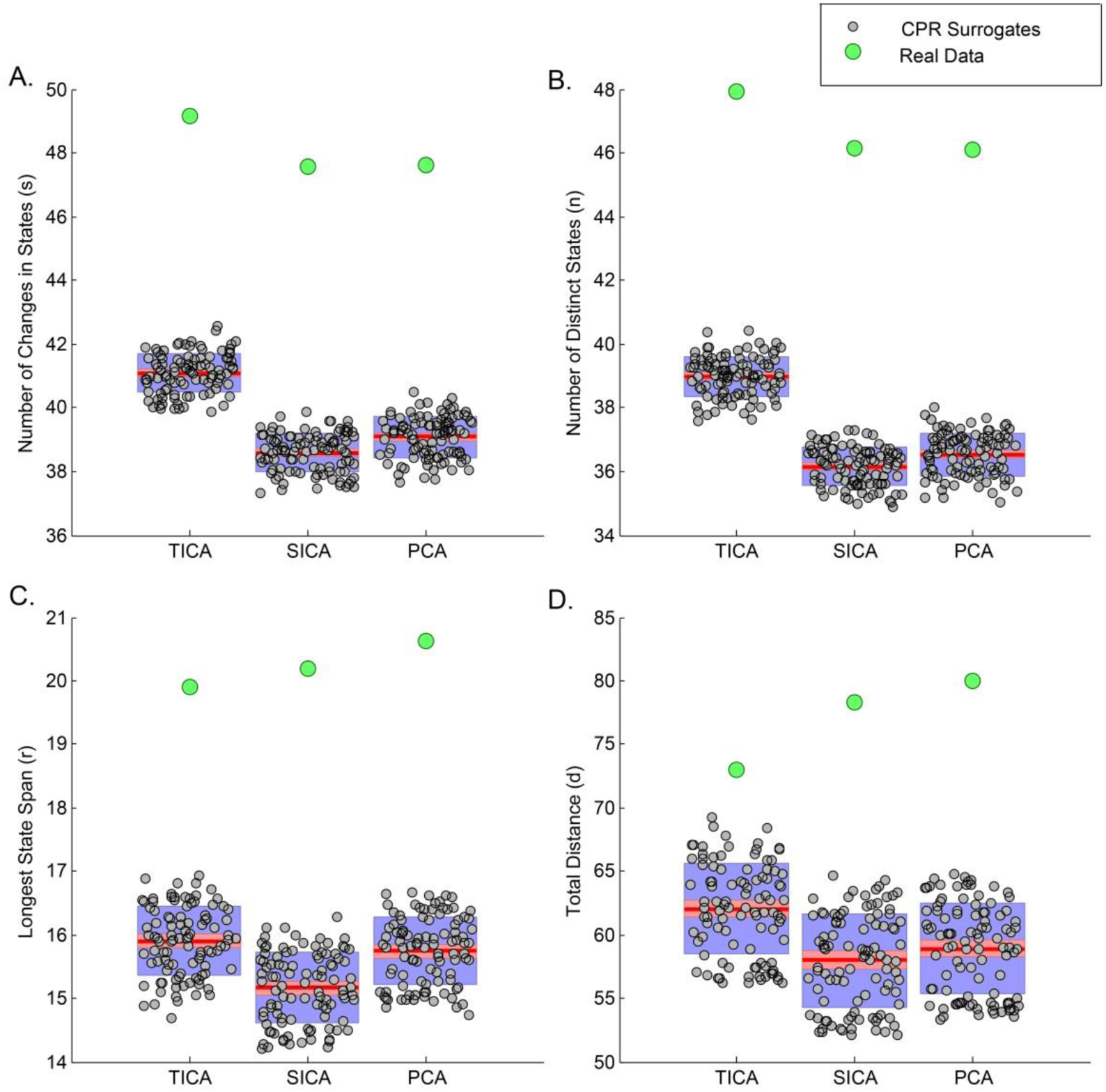
External validation in the meta-states approach. Meta-state metrics corresponding to real fMRI data were observed to fall outside the respective null distributions generated from meta-state metrics corresponding to 100 CPR surrogate datasets of RSN time-courses. Results for the temporal ICA, spatial ICA and PCA decomposition methods are shown; similar result was observed for k-means decomposition method.

## 4. DISCUSSION

Prior studies on assessment of spatiotemporal dynamic FC in the human brain have made use of similar dFNC approaches to characterize pathophysiology i.e. identification of disease states, thus corroborating the utility of the undertaken dFNC approaches (Damaraju, Allen et al. 2014, Rashid, Damaraju et al. 2014, Yu, Erhardt et al. 2015, Du, Pearlson et al. 2016, Miller, Yaesoubi et al. 2016). These studies found extensive additional information through use of these dynamic approaches as compared to that from static assessment of FC, hence advocating the use of dynamic analyses for better understanding of functional connectivities in the brain. Furthermore, the dFNC measures have been reported to relate to demographic characterization (Hutchison and Morton 2015, Yaesoubi, Allen et al. 2015, Yaesoubi, Miller et al. 2015, Preti, Bolton et al. 2016), consciousness levels (Hutchison, Womelsdorf et al. 2013, Amico, Gomez et al. 2014, Hudson, Calderon et al. 2014, Barttfeld, Uhrig et al. 2015, Wang, Ong et al. 2016) and cognition (Kucyi and Davis 2014, Schaefer, Margulies et al. 2014, Yang, Craddock et al. 2014, Madhyastha, Askren et al. 2015). Despite fundamental evidence of availability of considerable, interesting spatiotemporal dynamic connectivity information through these dFNC approaches, no prior study has yet evaluated the canonical utility of the dynamic measures; in other words, are there certain connectivity patterns that tend to recur across different subjects, i.e. a chronnectome.

To resolve this important issue, we tested two specific dFNC frameworks used in chronnectomic studies over multiple age-matched, large and independent resting state datasets. From group analysis in our first approach, we confirmed high correlation in sorted state profiles across the independent decompositions with the first quartiles (25^th^ percentiles) of pairwise correlations greater than 0.8 for 4 out of 5 states. We also observed consistent clustering results for a large range of clusters both of which clearly suggest faithful reflection of high degrees of replicability in the structure of underlying dynamics. Surrogate data analysis for this approach confirmed clustering to be substantially (but not completely) explained by the lagged cross-covariance structure of the RSN time-courses, while also suggesting the presence of non-linearity or non-stationarity, or both non-linearity and non-stationarity in this observed data. Using tSNE as a quality control measure, we successfully projected the high dimensional state profiles from all independent groups onto a two-dimensional space and could confirm existence of structure in the windowed FC data from all groups, and infer results consistent with the metrics derived in this approach. However, this visualization also suggested possible improvements in the chosen clustering algorithm since one of the states appeared as a mixture of two other primary states. Using the fuzzy meta-state approach as a second pass analysis, we evaluated multiple decomposition methods to explore generic replicability from a different perspective. As expected, we found low values of standard deviation for all derived average group-wise meta-state metrics through the temporal ICA decomposition method for a range of dimensions or model orders. This early identification was found to be consistent with similar evidence from replicability analysis using the k-means, spatial ICA and PCA decomposition techniques. Finally, validation analysis through permutation testing on the synthesized surrogate datasets confirmed evidence of presence of non-Gaussianty, non-linearity or non-stationarity (or any combination of these properties) in the observed RSN time-courses.

Head-motion has been shown to significantly alter correlations in the FNC (Power, Mitra et al. 2014), and so we studied the impact of head motion in the estimated replicability metrics. In this particular analysis, we de-spiked the data as well as regressed out the estimated head motion parameters from the observed dynamic measures, and derived the replicability metrics from these motion regressed dynamic measures. As seen in SI Figures 3 and 4, the regressed dynamic measures in both approaches were found to be very similar to the originally estimated dynamic measures and demonstrated similar replicability, suggesting head motion does not play a major role in the observed replicability.

Current findings from our analyses provide a substantial and novel advancement on the debate of robustness of inferences on temporal dynamics through the undertaken methods. Taken together, results from our analyses provide several lines of evidence of substantial reproducibility in the basic connectivity patterns amidst an ensemble of inter-regional connections, and further evidence of robust reproducibility across the independent decompositions against variation in data analysis methods, grouping methods, decomposition techniques and data quality. This evidence was found in a “probably” mixed dataset; however, we expect higher similarity in the dynamic FC measures in data with healthy controls only or patients only as could be expected to be obtained from analysis of a homogeneous sample. While our work confirms replicability of time-varying FC states (as evaluated using the two undertaken approaches) in the BOLD fMRI data, we still need to disambiguate variations in FC due to neuronal activity from variations due to other non-neuronal related phenomenon, for example, variations due to parametric choices in methodology or inherent noise in the BOLD fMRI data. The general points about the methodological considerations, parametric choices, need for generative null models to test for stationarity and neurological relevance of the observed dynamics as discussed exclusively in several recent studies (Lindquist, Xu et al. 2014, Zalesky, Fornito et al. 2014, Leonardi and Van De Ville 2015, Zalesky and Breakspear 2015, Hindriks, Adhikari et al. 2016, Miller and Calhoun 2017, Miller, Robinson et al. 2017) are certainly important.

The impact of parametric choices on the estimated dynamic measures or detection probabilities of the studied dynamic measures in the sliding window approach has been extensively studied (Lindquist, Xu et al. 2014, Leonardi and Van De Ville 2015, Zalesky and Breakspear 2015, Hindriks, Adhikari et al. 2016). Lindquist, Xu et al. (2014) focused on the problem of estimating temporal variation of bivariate correlations (pair-wise correlations between two fMRI time-series), comparing the sliding window (SW) method (for different window lengths), the tapered sliding window (tSW) method (for only one fixed window length), and two multivariate volatility models commonly used in finance literature, namely the exponential weighted moving average (EWMA) model and the dynamic conditional correlation (DCC) model. This analysis provides insights into general ability of longer window lengths for robust estimation of the correlation coefficients using the sliding window method as evident by overall performance and error measurements comparable to the claimed best method overall (DCC) in several tests conducted. Generally, a tapered sliding window is expected to produce smoother and more efficient correlation estimates than a rectangular window of the same window length and other window parameters, and hence, has been used in the majority of dFNC studies using the sliding window approach (see table S1 in Preti, Bolton et al. 2016 for a complete overview); however, from Lindquist, Xu et al. (2014) nothing much can be directly concluded in regards to performance of the tapered sliding window approach as only one configuration was compared to the other models. A more recent study (Choe, Nebel et al. 2017) compared the reliability of the mean and variance of the estimated network-pair correlations and FC state measures derived using the SW, tSW and DCC methods. This study concluded similar reliabilities in the estimated means across all estimated methods, improved reliability in the estimated variance for the DCC method (that advocates additional exploration of this alternate method in future FC investigations), and lower reliability of a few FC state measures (that is strikingly different to results in this paper but could be attributed to important methodological differences between the two works, for example, initialization of the clusters with exemplar data which was done in our work as well as Allen et al, but not used in Choe et al 2017). Studying dFNC simultaneously with multiple proven methods and focusing on overlapping, more consistent results would enhance reliability of the inferences. Furthermore, Leonardi and Van De Ville (2015) highlighted the importance of systematic study and practical guidelines for the choice and impact of different chosen parameters in a sliding window approach. The authors recommended the “1/f rule of thumb” for calculating minimal window length while observing underlying frequencies in the fMRI signal. Zalesky and Breakspear (2015) reviewed these findings providing statistical support for their proposed rule of thumb, but additionally making a point that the proposed lower limit is overly conservative especially in moderate SNR conditions. The latter findings are corroborated in several studies where varying the window length parameter over a range beyond a certain safety limit did not change the overall observed dynamics (Allen, Damaraju et al. 2012, Li, Zhu et al. 2014, Yaesoubi, Miller et al. 2015, Deng, Sun et al. 2016, Liégeois, Ziegler et al. 2016, Preti, Bolton et al. 2016).

Hindriks, Adhikari et al. (2016) evaluated the ability of the sliding-window correlations in revealing dFNC and also highlighted the importance of appropriate statistical tests to detect dFNC. In this work, the authors set up a model for the dynamics of the FC time-series using a controlled “correlation timescale” (ҭ), “dFNC strength” (η), scan length, sliding-window length and sliding-window step-size parameters, and use it to quantify the ability of a linear (standard deviation of the windowed FC data) and a non-linear (as proposed in Zalesky et al. 2014) test statistic to detect dFNC by estimating a formally defined, statistical measure, namely “detection probability”, for a range of the controlled parameters. While this particular analysis also suggested higher detection probabilities for increased values of both “ҭ” and “η” in the simulated data, one of the major conclusions drawn from this study, namely the inability of the sliding-window method to detect dFNC, is based on a particular experiment wherein the estimated detection probability was low while assuming fixed values for correlation timescale and dFNC strength parameters. Both of these assumed parameters are unknown for real fMRI data. Clearly, the observations are not unconditional because of their dependence on these unknown parameters, and hence this particular conclusion from their experiment cannot be fully generalized to real fMRI data. Additionally, in another part of this analysis, the authors mention the absence of any information regarding how to set these two parameters and estimate the optimal value of the sliding-window length parameter by averaging the observed detection probabilities over all values of correlation timescales (ҭ). They conclude the optimal value of sliding-window length strikingly close to the commonly chosen duration of 60 seconds. Our work and most of the studies using the sliding window approach use a similar duration for the sliding-window, a point also discussed in major reviews in the area (Calhoun, Miller et al. 2014, Calhoun and Adali 2016, Preti, Bolton et al. 2016). Subsidiary methodological variations must also be carefully tested for any major effects; for example, clustering the PCA reduced windowed FNC data (reduction on the ROI-pair correlation dimension as in Laumann, Snyder et al. (2016) retains a smaller percentage of variability in the data and tends to drive apparent similarity (reduce distance) in the emergent state profiles (demonstrated in SI Figure 2). As a note, despite overall consistency, the methodology steps and other parametric choices in the sliding-window method must continually both theoretically and empirically be explored to further improve accuracy of the inferences.

Evaluating statistical significance of the estimated dFNC measures assumes similar importance as making appropriate methodological and parametric choices. Undoubtedly, it would be highly useful to replicate the behavior of “noiseless” BOLD data by “appropriate” simulations; however, an absence of a baseline, i.e. ground truth for resting state, makes this very step extremely challenging. Null models must therefore be approximated using the available fMRI data. Previous research has used the CPR and vector auto-regressive (VAR) models for this purpose. These models allow testing for the hypothesis that the observed data is generated by a linear, stationary Gaussian process. In this study, we saw the CPR null model being rejected for one test statistic in the hard clustering approach and four test statistics in the meta-state approach. A major drawback of this null hypothesis is that it too general, and if rejected, it is not possible to conclude the specific cause of rejection to be non-Gaussianity, non-linearity, non-stationarity or some combination of these properties, and hence there is need for additional analysis. In case of rejection of these null models, it would make sense to test for Gaussianity of the observed data as it is more straightforward, and if the data is concluded to be Gaussian, subsequent advanced statistical testing, for example testing the degrees of non-stationarity and non-linearity, could be explored to further comment on the specific property causing the rejection of the null model. Other recently used alternatives to the CPR and VAR null models include the amplitude-adjusted phase randomization (AAPR) null (Betzel, Fukushima et al. 2016) and the null as used in Laumann, Snyder et al. (2016). The null hypothesis in AAPR model associates to the observed data being a monotonic non-linear transformation of a linear Gaussian process (Theiler, Eubank et al. 1992, Schreiber and Schmitz 2000) and generates data that preserves the amplitude distribution exactly but the power spectrum approximately. Finally, the null used in Laumann, Snyder et al. (2016) is matched to the covariance structure exactly i.e. preserves the static FC exactly, but to the power spectrum on average i.e. does not preserve the cross-lagged covariance structure exactly as in the CPR and VAR models. Since these different models correspond to different null hypothesis and preserve different properties of the observed data, exploring and utilizing additional knowledge on the nature of the observed data is recommended to appropriately choose the null hypothesis in a given study. In nutshell, there is need for additional work in the field of null model development for statistical validation by surrogate testing, and hopefully more specific null models and/or frameworks to test existing null models in literature will emerge and eventually allow for more specific inferences.

Besides, some innovative ways of using null data to draw conclusions about time-varying nature and consistency of the FC fluctuations have also been recently explored (Zalesky, Fornito et al. 2014, Betzel, Fukushima et al. 2016, Hindriks, Adhikari et al. 2016). Zalesky, Fornito et al. (2014) used a novel framework to provide evidence of a consistent set of “dynamic” inter-RSN connections that exhibited pronounced fluctuations in strength over time. This framework records a non-linear “excursion” test statistic quantifying the extent of time-varying fluctuations in the windowed FNC data for both original data as well as a set of VAR surrogate datasets. In the next step, connections that reject the null distributions of this test statistic are retained for further analysis wherein binary graphs are constructed for each subject using only the top-few most “dynamic” connections and degree of each region in these binary graphs is evaluated. Finally, this degree is summed across the subjects to frame an “index of consistency” of these dynamic connections (i.e. how consistently the regions were dynamic across the subjects). Next, Hindriks, Adhikari et al. (2016) reported absence of evidence for dFNC in real fMRI data for individual sessions and concluded that using the CPR null model it is difficult to distinguish two test statistics, namely the standard deviation of the windowed data and the non-linear test statistic as originally explored in Zalesky, Fornito et al. (2014). Our analysis with the CPR surrogate datasets generated from fMRI data used in our study suggests the presence of significant “dynamic” inter-regional connections which we also evaluated for consistency through the “index of consistency” metric for both the standard deviation of windowed FNC data and the excursion test statistics (SI Figure 1). Evaluating consistency of these significantly “dynamic” inter-regional connections across numerous independent samples similar to this study is definitely an interesting work for future.

Recently, Laumann, Snyder et al. (2016) suggested stability of the FC structure observed in resting state BOLD fMRI data over tens of seconds. The authors clearly mention in their work that this demonstrated stability of the FC structure, computed by integrating over time, did not cross paths with time-varying studies analyzing shorter time-scales. Furthermore, recent collaborative work from the same authors has formally demonstrated that even statistically stationary data does not imply absence of brain states (Liegeois, Laumann et al. 2017). The authors also suggested the emergence of the observed states mostly due to sampling variability and physiological confounds in the fMRI data. Our perspective on sampling variability is that such variability (between subjects) is certainly possible but does not by itself argue for or against the presence of dynamic states any more than sampling variability visible in an analysis of GLM maps enables us to detect the presence of the widely studied resting networks in second level task-based fMRI data (Smith, Fox et al. 2009, Allen and Calhoun 2012) argues against the presence of resting fMRI networks. In addition, there does appear to be agreement that the FC fluctuations in the data ‘indistinguishable’ from statistically stationary null data could demonstrate behavioral relevance (Shine and Poldrack) and electrophysiological correlates as discussed in the concluding paragraph.

The argument whether the functional connectivity data is better modeled by “states” or “meta-states” as by frameworks used in this paper or with other proposed dynamic models (for example, an autoregressive model) is still a matter of debate. It must also be noted that there could be several different ways of capturing the temporal dynamics in fMRI data and in turn illuminating the brain function; the methods studied in this paper do not claim a specific number of states in the fMRI data any more than a specific number of resting state networks in fMRI data could be claimed. Rather, the focus is on illustrating that such a decomposition may be useful for studying the brain, and this necessitates the ability to identify stable connectivity patterns from the data that can replicate and which show similar temporal dynamic properties. There is already evidence for usefulness of the dynamic state models explored in this work as previous work has demonstrated that such patterns are better than static connectivity at predicting patient groups which suggests that such decompositions as explored may be useful for helping differentiate patients and controls (Damaraju, Allen et al. 2014, Rashid, Damaraju et al. 2014, Yu, Erhardt et al. 2015, Du, Pearlson et al. 2016, Miller, Yaesoubi et al. 2016).

Going forward, investigating the functional and neurophysiological relevance of the observed time-varying FC states, meta-states or other robust connectivity descriptors assumes critical importance and needs further confirmation. Demonstrating functional relevance is currently an active topic with several interesting works establishing direct links with ongoing cognitive function and effective cognitive performance (Craddock, James et al. 2012, Schaefer, Margulies et al. 2014, Gonzalez-Castillo, Hoy et al. 2015, Madhyastha, Askren et al. 2015, Shine, Bissett et al. 2016, Shine, Koyejo et al. 2016), identifying signatures of consciousness (Hutchison, Womelsdorf et al. 2013, Amico, Gomez et al. 2014, Hudson, Calderon et al. 2014, Barttfeld, Uhrig et al. 2015, Wang, Ong et al. 2016), tracking day-dreaming/mind-wandering (Kucyi and Davis 2014, Kucyi 2017), and decoding signatures of sleep and awake states (Tagliazucchi and Laufs 2014). Additionally, simultaneous recording of electrophysiological data in conjunction with BOLD fMRI data not only enables charting of the human brain activity at high spatial as well as high temporal resolutions, but also linking variability in FC fluctuations to external measures of neuronal activity. Recently found evidence of potential electrophysiological signatures of dynamic BOLD FC clearly hint fluctuations in the BOLD FC to be interesting i.e. having a neurophysiological origin (Tagliazucchi, von Wegner et al. 2012, Chang, Liu et al. 2013, Allen, Damaraju et al. 2017). Such preliminary observations clearly suggest that multi-modal studies may play a key role in determining neural or behavioral relevance of the observed FC states in the fMRI data. Future work would hence likely involve recognition of a multi-modal, multi-level theoretical framework that would very likely be able to capture the underlying physiological correspondences that enable switching of the emergent connectivity patterns. Neuronal correlates of time-varying FC have been previously suggested in few studies (Liu, Chang et al. 2013, Thompson, Merritt et al. 2013, Kragel, Knodt et al. 2016), and more recently Matsui, Murakami et al. (2017) claims a link between time-varying FC and neuronal origins, however, a lot of additional work is still needed to affirm the correspondence of the time-varying FC state descriptions with underlying neuronal activity. Other lesser explored alternate approaches that have shown promise include casual manipulation of the FC states using pharmacology (Hutchison, Womelsdorf et al. 2013, Barttfeld, Uhrig et al. 2015, van den Brink, Pfeffer et al. 2016) or direct brain simulation techniques (Liu, Lee et al. 2015, Cocchi, Sale et al. 2016) to trace functional or neurophysiological relevance (Shine and Poldrack). We predict such tailored investigations probing the functional and neurophysiological relevance of the time-varying FC states will play a major part in uncovering the underlying relationships we are pursuing; replicability is only an initial step forward.

## ACKNOWLEDGEMENTS

The authors thank Charlotte Chaze for her contribution to the group’s 2016 EMBC paper, the results of which motivated further research on this project, Srinivas Rachakonda for lending his expertise on the GIFT toolbox functions, Helen Petropoulos for providing information on fMRI data analyzed in this paper and Dr. Maziar Yaesoubi for his advice on data analysis methods. The authors declare no competing financial interests.

